# From constraint to opportunity: Relaxing sexual antagonism reveals adaptive potential maintained by balancing selection

**DOI:** 10.64898/2026.02.04.703537

**Authors:** Martyna K. Zwoinska, R. Axel W. Wiberg, Milena Trabert, Philipp Kaufmann, Elina Immonen

## Abstract

Despite ongoing selection, genetic variation in fitness-related traits often persists. Balancing selection can maintain polymorphisms through genetic trade-offs, including those between the sexes. Sexually antagonistic (SA) selection is challenging to detect at the genomic level and its broader evolutionary importance, especially for complex traits, remains unclear. To investigate this, we conducted an evolve-and-resequence experiment in *Callosobruchus maculatus*, manipulating the strength of SA trade-offs over body size and tracking genome-wide responses. When selection simultaneously favored larger females and smaller males, allele frequency changes were constrained and genome-wide divergence remained limited. In contrast, relaxing SA trade-offs by selecting on only one sex led to large, repeatable allele frequency shifts. These loci also showed signatures of long-term balancing selection in the ancestral population. Together, our results demonstrate that SA trade-offs can act both as a constraint, limiting sex-specific responses under antagonistic selection pressures, but also as a source of adaptive potential once antagonism is relaxed.

## Introduction

The persistence of genetic variation in the face of selection and how such variation contributes to adaptive potential remains a central question in evolutionary biology ^1^. Balancing selection, an umbrella term for selective scenarios that maintain genetic polymorphisms, is often invoked to explain high levels of genetic variation observed in fitness-related traits ^2–5^. Yet its evolutionary importance remains unresolved, a debate that has persisted for over a century and has recently been reignited by the influx of genomic data ^6–8^.

Sexually antagonistic (SA) selection is a well-recognized scenario that can give rise to balancing selection ^9–15^. SA selection occurs when trait optima diverge between the sexes due to differences in reproduction, ecology, or life history ^16–18^. Because most genes are shared between the sexes, selection on divergent trait optima can cause the same allele to be beneficial in one sex but costly in the other, resulting in SA genetic trade-offs ^12,17,19^. Although genetic trade-offs can also arise when selection coefficients differ in sign across space or time, SA trade-offs are unique in that antagonism is intrinsic and simultaneous: opposing selection stems from the concurrent demands of both sexes acting on a shared genome, preventing either sex from reaching its fitness optimum. As a result, alleles with sex-opposite fitness effects may be maintained in the population instead of fixing or being lost.

Theoretical models have extensively explored the conditions under which SA trade-offs can maintain genetic polymorphisms. Single-locus models support the persistence of SA polymorphisms across a broad range of parameter values, particularly when selection is strong and symmetric, and when allelic dominance depends on the sex in which alleles are expressed ^9,11,13,20–23^. In contrast, polygenic models predict more restrictive conditions for maintaining SA variation, especially when a trait is controlled by many loci with small effects ^14,15^. However, it remains unclear how often natural populations meet these theoretical conditions, or whether the assumptions of the models adequately capture biological reality. Consequently, although studies have identified loci with opposing fitness effects between the sexes ^24–27^, the overall contribution of sexual antagonism to maintaining genetic variation remains uncertain.

While the focus has been on how genetic variation is maintained under SA trade-offs, it is equally important to understand whether and how this variation contributes to adaptation when trade-offs are relaxed. Such relaxation can occur under environmental change that aligns selection pressures between the sexes ^28,29^. In these cases, alleles previously maintained by SA trade-offs may fuel rapid adaptation, particularly if they have large effects that allow swift shifts across fitness landscapes ^30^. A few compelling examples exist in salmon and *Drosophila*, where large-effect SA variants appear to have facilitated adaptation to anthropogenic pressures ^31–35^. Adaptive genetic architecture of traits is however difficult to predict ^36^ and it remains unclear whether variation maintained due to sexual antagonism can meaningfully contribute to adaptive responses.

Here, we empirically address two key gaps in understanding the role of SA selection in adaptation: first, how SA trade-offs over a polygenic trait maintain genetic polymorphism^37^; and second, how such variation contributes to the genomic signal of adaptation when antagonism is relaxed. If SA trade-offs maintain alleles with opposing fitness effects, then relaxing this antagonism should release standing genetic variation from constraint and drive allele frequency divergence. In contrast, intensifying these trade-offs should limit genomic change. Moreover, if such antagonism has persisted over evolutionary timescales, loci that respond when it is relaxed may also carry signatures of long-term balancing selection.

To test these predictions, we used an experimental evolve-and-resequence approach in *Callosobruchus maculatus* seed beetles, tracking genomic changes during artificial selection on body size across replicated lines ^38,39^. Body size is a complex and often sexually dimorphic trait ^40^, with high heritability and a largely shared genetic architecture between the sexes in our study population ^38^. Our experimental design included several treatments. In the SA treatment, we intensified conflict by selecting simultaneously for larger females and smaller males within families. In three sex-limited (SL) treatments, selection targeted only one sex at a time, favouring either larger females, smaller males, or larger males, which reduced the level of genetic conflict. A Relaxed selection control line was maintained to benchmark genetic drift. We tracked allele frequency changes at the whole-genome level at the beginning, midpoint, and end of the experiment. Consistent with our predictions, as well as phenotypic responses (Fig. 1) ^38^ and pedigree data in the lines ^37^, we found widespread and directional allele frequency change when the SA trade-off was relaxed, but little changes relative to control when it was intensified. Our results highlight the dual role of sexual antagonism in both preserving genetic variation and shaping adaptive potential.

**Figure 1.**
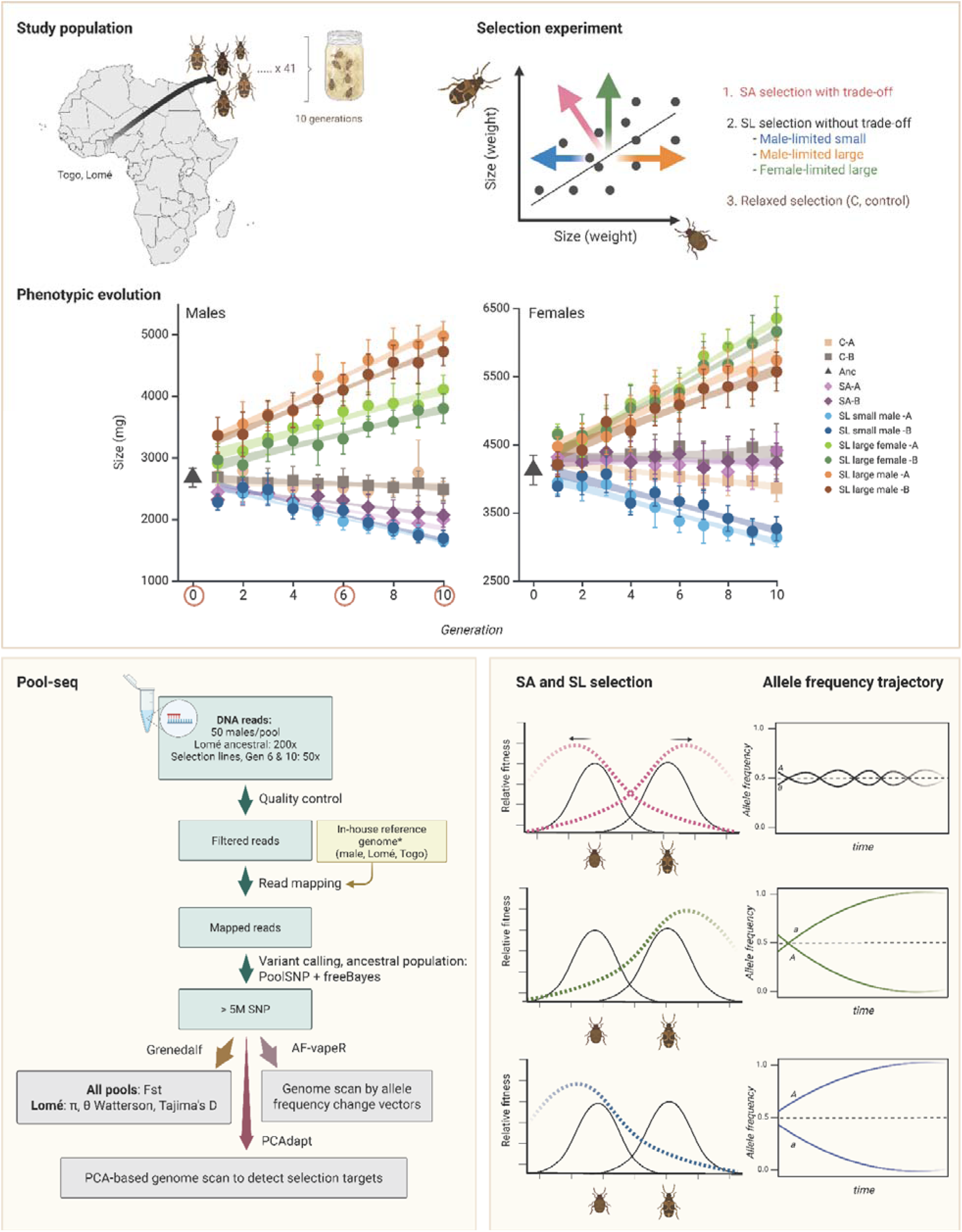
Experimental design. Upper panel: A wild-caught population of *C. maculatus* from Lomé, Togo, was used as the study population, established as isofemales lines that were subsequently joined into a single population 10 generations prior to selection. The selection treatments (applied at family-level) included intensification of SA trade-offs by selection for larger females and smaller males (SA), relaxation of SA trade-offs by limiting selection on one sex (SL), and relaxation of selection altogether (C, Relaxed selection). Phenotypic divergence was rapid and repeatable under SL but constrained under SA selection, especially female size. Lower panel: Samples for Pool-seq were collected from the ancestral population at parental generation to the selection lines, and at generations 6 and 10 of selection. SA selection is predicted to maintain allelic polymorphism at loci subject to divergent SL selection.

## Results

### Genomic patterns of divergence reveal constraint imposed by SA selection

Our analyses are based on 5.4 million high-confidence SNPs segregating in the ancestral population, that is, at the parental generation of the selection lines. Variants were identified through variant calling in this population (see Methods), and their frequency changes were tracked across all selection lines and generations. In all analyses, allele frequencies are expressed with respect to the ancestrally minor allele.

To examine the genomic consequences of both relaxing and intensifying SA selection, we first performed a principal component analysis (PCA) on autosomal SNP frequency data (Fig. 2, S1). The first principal component (PC1), which explains 12% of the total genetic variation, mirrors the phenotypic divergence observed between selection regimes (Fig. 1). SL lines, which were selected for large or small size in only one sex, fall at opposite extremes of PC1. This pattern demonstrates their rapid and divergent evolutionary responses when the SA trade-off is relaxed. The ancestral population, as expected, occupies a central position along PC1 and other major axes. The distance between ancestral and derived populations increases over time, from generation 6 to generation 10 (Fig. 2). Genome-wide pairwise *F_ST_* values across all autosomal SNPs support the PCA results, revealing a consistent increase in genetic divergence, from 0.039–0.047 at generation 6 to 0.055–0.084 at generation 10 (Fig. S2). SA lines, which evolved under intensified sexual antagonism, cluster closest to the ancestral population along PC1 and PC2. Similarly, lines under relaxed selection also remain genetically close to the ancestral population on PC1.

**Figure 2.**
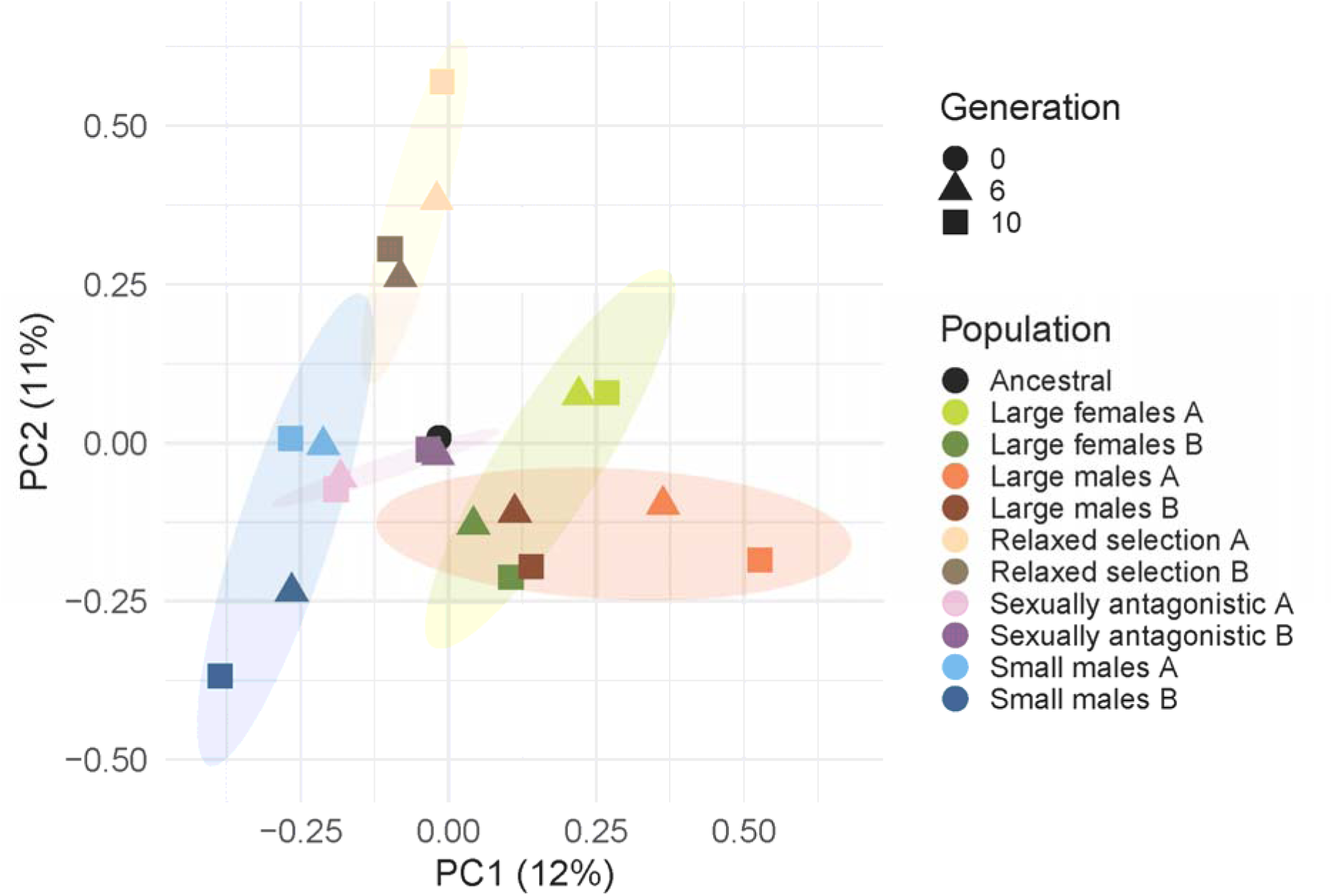
PCA plot of SNP frequencies at autosomal loci showing divergence of the selection lines from the ancestral population (generation 0, black dot), at generation 6 (triangles) and generation 10 (squares).

**Figure 3a.**
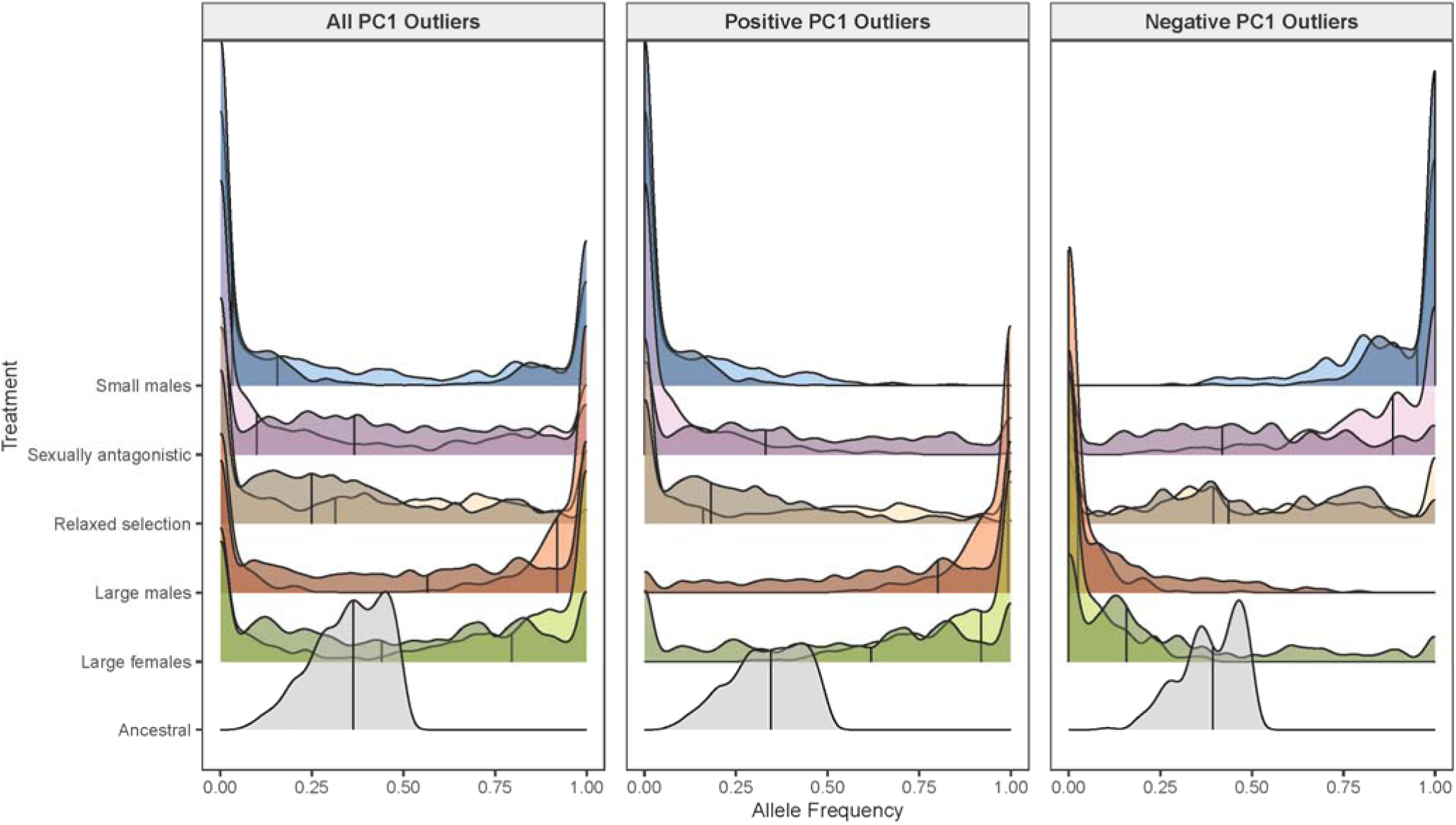
Allele frequency distributions for autosomal SNPs with significant loadings onto PC1, in each selection regime after selection (at generation 10) relative to the ancestral population (at generation 0). Replicate lines for each selection regime are shown with different shades. Vertical lines indicate the median frequency (relative to the ancestrally minor allele).

**Figure 3b.**
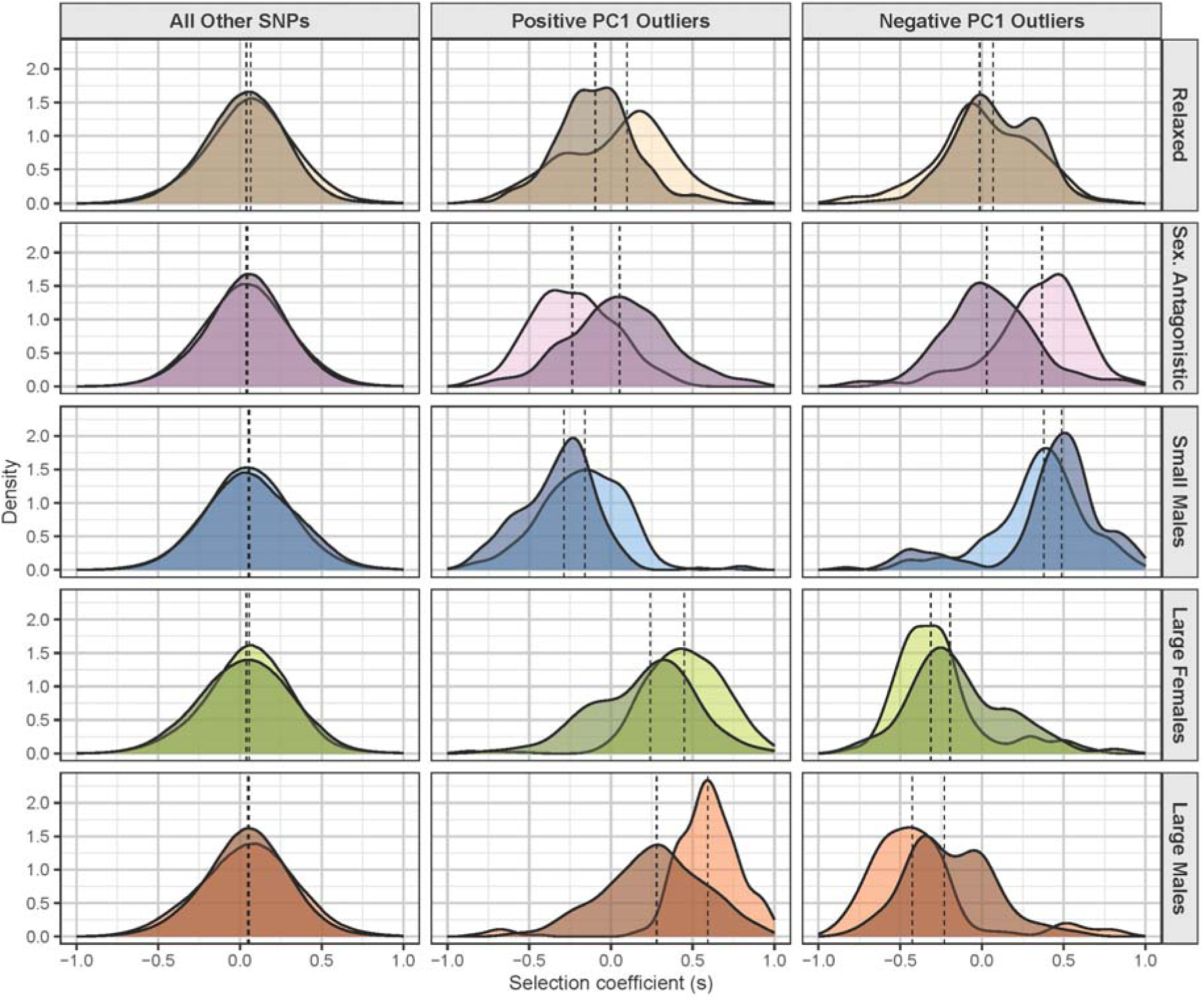
Selection coefficients (*s*) for autosomal SNPs with significant loadings onto PC1, in each selection regime after 10 generations of selection. Replicate selection lines are shown with different shades. Vertical lines indicate the median *s*.

**Figure 3c.**
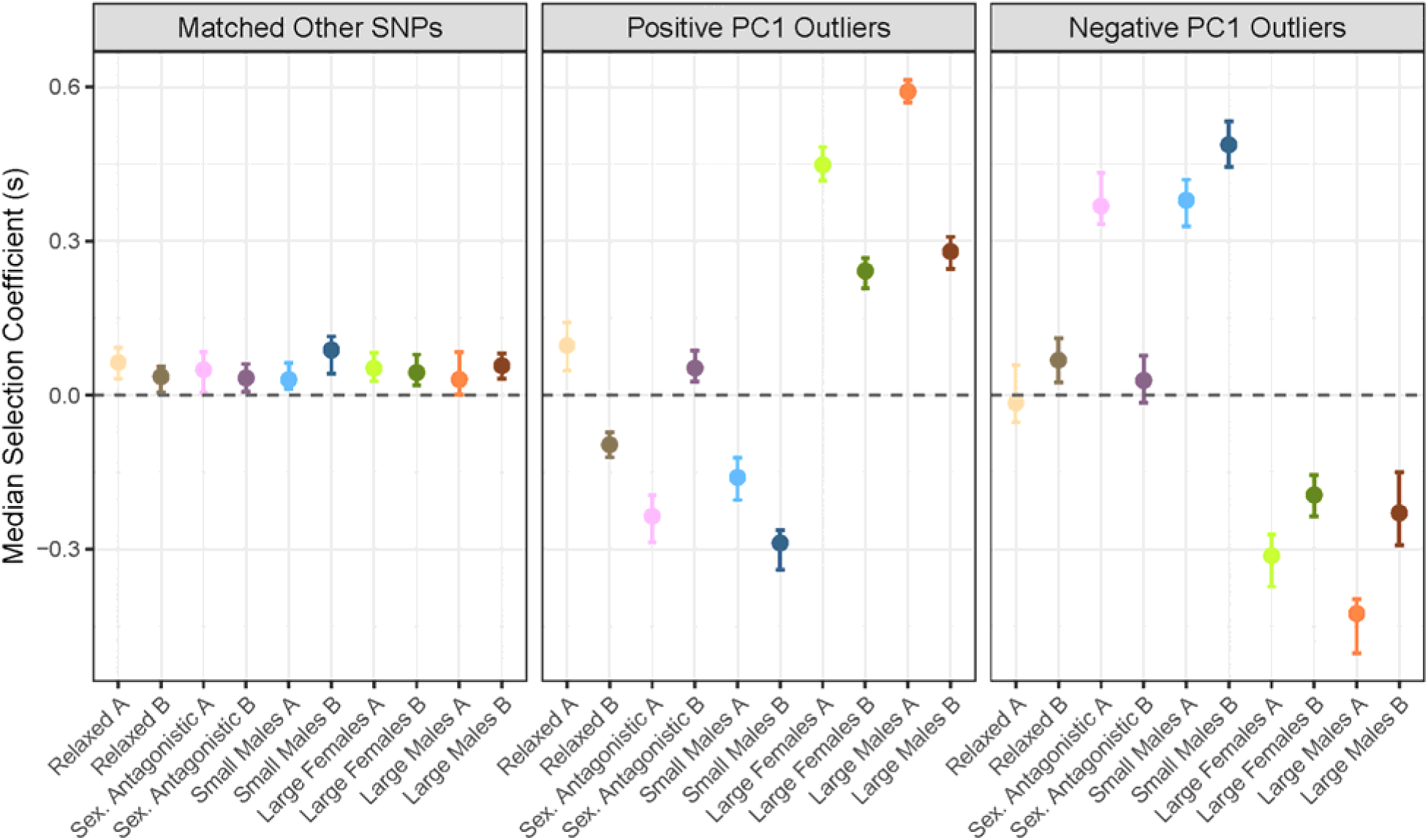
Median selection coefficients (*s*) (with bootstrapped 95% CIs) for autosomal SNPs with significant positive (“Positive PC1 Outliers”) or negative (“Negative PC1 Outliers”) loadings onto PC1, shown for each selection line. For comparison, an equal number of non-outlier SNPs (“Matched Other SNPs”) were randomly sampled within each selection line.

**Figure 4a.**
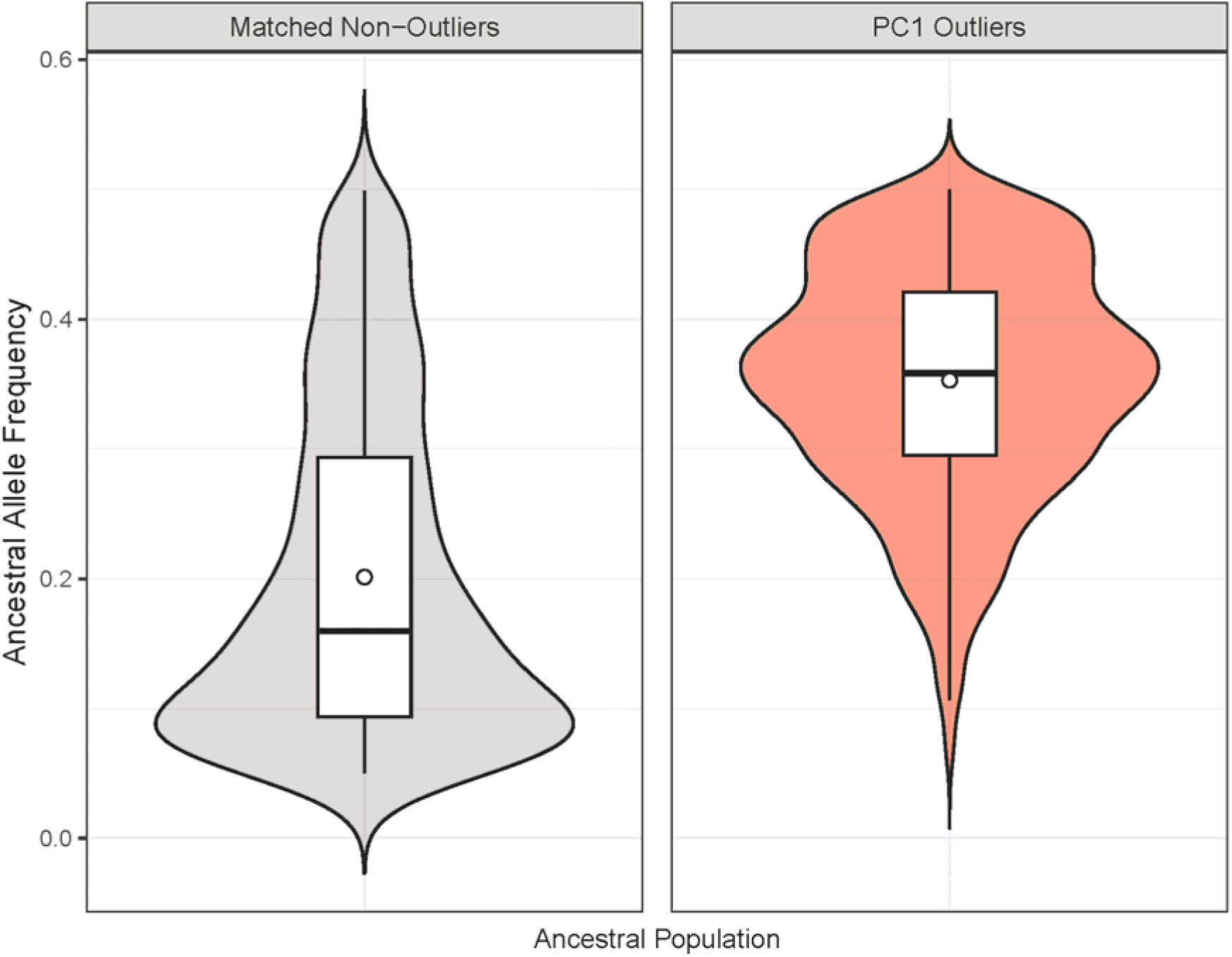
Outlier SNPs had higher ancestral allele frequencies than non-outliers in the ancestral population of selection lines. Non-outliers were randomly sampled within each selection line to match the number of outliers.

**Figure 4b.**
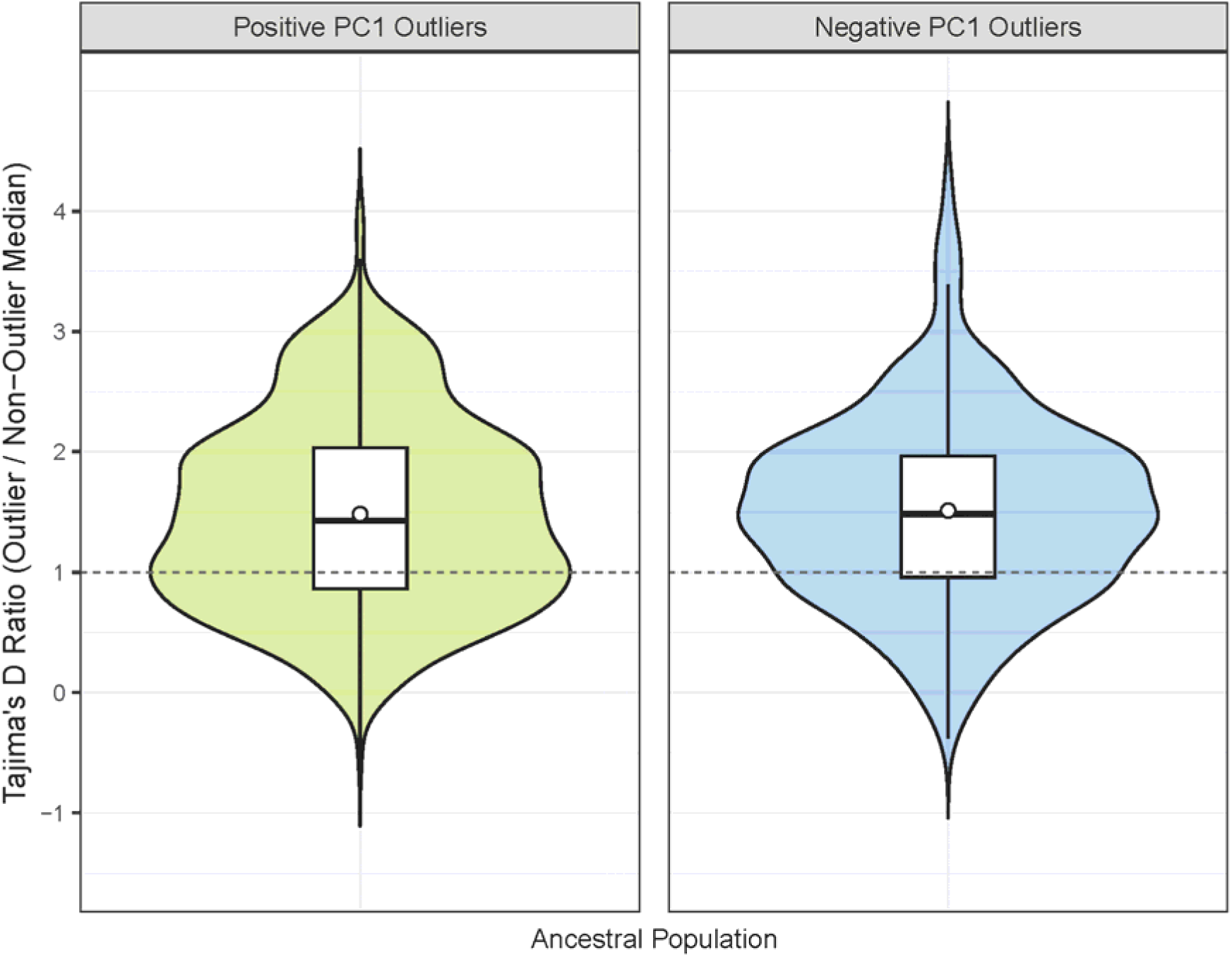
Tajima’s D values are elevated for PC1 outlier SNPs in comparison to non-outlier SNPs in the ancestral population.

PCA results confirm that the major axes of genomic variation align with the predicted evolutionary responses to selection. To complement this approach, we also quantified allele frequency shifts along predefined contrasts using AF-vapeR ^41^. Specifically, we defined vectors of allele frequency change from the ancestral population to each derived line at generation 10 and focused on genomic windows showing the strongest divergence between SL lines selected for small male body size versus those selected for large male or female size. Eigenvector loadings from these windows showed that SA and Relaxed selection lines clustered near the center of the divergence axis (Fig. S3), consistent with constrained genomic responses under SA selection, as also observed in the PCA.

We next examined divergence on the sex chromosomes to determine whether patterns mirrored those on the autosomes. PCA of Y-linked SNPs revealed a distinct pattern: SL lines selected for large male size separated clearly from all other treatments along PC1, which accounted for 40% of the total variance (Fig. S4d). *F_ST_* values confirm the elevated divergence of SL Large males (Y-linked *F_ST_* range between Large male lines and the ancestral population and Small male lines, respectively: 0.136–0.138; 0.197–0.258 Fig. S5b). These Y-linked patterns are consistent with the presence of a previously identified, functionally distinct Y haplotype in the Large male lines ^39^. The X chromosome also shows some degree of structure, but unlike the Y and the autosomes, it does not reflect the selection history (Fig. S4c, S5a).

### Relaxation of the SA trade-off leads to rapid and consistent allele frequency divergence

Next, we sought to identify loci underlying the evolved population structure revealed by PCA. We applied a component-wise genome scan using the pcadapt package ^42^ to detect SNPs whose allele frequency variation across populations is significantly associated with individual principal components. This analysis identified 869 SNPs significantly associated with PC1 (q < 0.05, Fig. S6). These outliers are widely distributed across the genome, indicating that the observed divergence reflects polygenic responses to selection (Fig. S7, 3b). We then examined how allele frequencies at these PC1 outlier loci changed across selection regimes. Because PCA loadings reflect the direction of association between SNPs and population structure, SNPs with opposite PC1 loadings should exhibit opposing allele frequency changes across treatments. Consistent with this expectation, SNPs with negative PC1 loadings increase in frequency under selection for smaller body size and decrease when selection favored larger size, whereas SNPs with positive PC1 loadings show the opposite pattern (Fig. 3a). Together, these patterns reveal a consistent, trait-associated genomic response to selection.

Having identified PC1 outlier loci underlying the evolved population structure, we quantified divergence from the ancestral state at these loci as the mean absolute allele frequency change (|ΔAF|) from generation 0 to generation 10, and compared this measure across treatments. Divergence under SA selection was much closer to that observed under relaxed selection than under SL Small male selection (paired Wilcoxon signed-rank tests on |ΔAF|: SA vs Relaxed, W = 674 964, p < 10^-10^; SA vs Small males, W = 344 585, p < 10^-23^).

Finally, we estimated selection coefficients (*s*) for segregating variants using our time-series allele frequency data, using the poolSeq package ^43^. We focused on two internal comparisons: (1) across selection regimes, using the Relaxed selection lines as a baseline reflecting primarily drift; and (2) between PC1 outliers (identified via pcadapt) and the genomic background (i.e. non-outliers). Selection coefficients aligned with SNP loadings: SNPs with negative PC1 loadings (“negative PC1 outliers”) showed on average a high positive *s* under SL selection for small size and a negative *s* under SL selection for large size, regardless of the targeted sex. The opposite pattern was observed for positively loaded SNPs (Fig. 3b-c). SL line replicates showed consistent and concordant selection: both replicates exhibited the same direction of *s*, and average values were significantly different from zero. In contrast, Relaxed and SA selection replicates showed more variable responses, including opposing directions of *s* across replicates (Fig. 3b-c). Together, these results indicate that relaxing SA trade-offs via SL selection produced the most pronounced selection responses. In contrast, intensifying SA selection led to weaker and more variable patterns, resembling those observed in the Relaxed selection lines.

### Loci targeted by SL selection show signatures of balancing selection

Having shown that sexual antagonism constrains short-term genomic responses by impeding fixation at loci associated with body size variation, we next asked whether these loci are more polymorphic in the ancestral population. In support, we found that PC1 outliers responding to SL selection segregated at higher ancestral minor-allele frequencies than non-outliers (median MAF: 0.36 vs. 0.16; 95% CI: 0.355–0.372 for outliers, 0.159–0.160 for non-outliers; Fig. 4a). Outliers were also depleted among less common variants (MAF 0.05–0.1: 0.006 for outliers vs. 0.29 for non-outliers) and enriched among intermediate-frequency variants (MAF 0.3–0.4: 0.335 vs. 0.125; 0.4–0.5: 0.363 vs. 0.109, outliers vs. non-outliers), a pattern consistent with balancing selection. Although outlier detection methods such as pcadapt are more sensitive to variants with intermediate starting frequencies, the observed enrichment aligns with PC1 divergence between Large and Small SL lines (Fig. 2), indicating that the signal likely reflects polygenic selection rather than a detection artefact.

To further test the hypothesis of balancing selection, we examined Tajima’s D (*T_D_*), a statistic that increases when there is an excess of intermediate-frequency variants. We first compared *T_D_* values between PC1 outlier windows and all non-outlier windows. On average, outlier windows had significantly higher *T_D_*, consistent with elevated sequence diversity (t_(861)_ = 18.6, p < 2.2×10^-16^; Fig. 4b). To control for variation in coverage and SNP density, we then repeated the comparison using matched non-outlier windows (with similar SNP counts and coverage) across three levels of outlier density: Low, Medium, and High (windows ≥ 20 SNPs). For each stratum, we calculated Δ, the mean paired difference in *T_D_* (outlier − matched). *T_D_* differences were smallest but still significant in the Low stratum (n = 153 pairs; Δ = 0.161; 95% CI [0.029, 0.294]; permutation p = 0.0136), increased in Medium (n = 146; Δ = 0.277; 95% CI [0.130, 0.423]; p = 0.001), and were largest in High (n = 149; Δ = 0.398; 95% CI [0.246, 0.549]; p = 0.0002). These results indicate that regions enriched for PC1 outliers exhibit systematically higher *T_D_*, with effect size increasing with outlier density.

Collectively, this suggests that the regions most responsive to SL selection, where sexual antagonism was relaxed, harboured pre-existing polymorphisms with allele frequency spectra and sequence-level diversity patterns consistent with long-term balancing selection.

### Gene Ontology analysis identifies divergence in BMP signaling in absence of SA trade-off

The majority (77%) of the PC1 outlier SNPs are intergenic and within 50kb distance from coding regions. 211 SNP’s (24% of all outlier SNPs) are in approximate regulatory regions of genes (i.e. SNPs within 2kb up/downstream from genes), while 10 SNPs reside inside the coding regions of 10 genes (Table S1). There are 72 genes with SNPs under selection for larger size (i.e. SNPs significantly loading to the positive side of PC1), while 48 genes have SNPs under selection for smaller size. GO analyses (Biological Processes) point towards divergence in early development processes, including BMP signaling, muscle tissue, neuron development, and cardiac cell fate specification during gastrulation (Table S1). Importantly, 11 genes have SNPs associated with evolution in both directions, 24% of all the implicated genes (Table S1). One of these genes under divergent selection in SL lines is a *C. maculatus* ortholog of *Tolloid* (*Tld*), a conserved growth regulatory member of the BMP pathway, which was independently identified using both the unsupervised and supervised outlier detection approaches. *Tld* allele frequency changes exemplify the predicted pattern where intermediate frequency SNPs in the ancestral population respond strongly and consistently to bidirectional selection when SA trade-off is relaxed, but not when it is intensified nor when body size is under relaxed selection (Fig. S8).

## Discussion

Genetic trade-offs have long been recognised as a potential source of balancing selection, and recent work suggests they may play a more widespread role in maintaining genetic variation than previously appreciated. These trade-offs arise when selection favours different alleles in different contexts, including between sexes, environments, or fitness components. Although an increasing number of SA loci have been identified, the overall genomic consequences of SA selection and its importance for both short-term change and the long-term maintenance of polymorphism remain poorly understood, particularly for complex traits. To address this, we combined replicated artificial selection on body size with genomic time series data. This allowed us to test how alleles constrained by SA trade-offs respond when conflict is relaxed, and whether they carry signatures of balancing selection. We find that loci maintained at intermediate frequencies responded rapidly under sex-limited selection and also exhibit hallmarks of long-term balancing selection, suggesting that SA trade-offs can help preserve genetic variation even in complex traits like body size.

Our first major finding comes from analysing short-term genomic responses to artificial selection under contrasting sexual selection regimes, where SA trade-offs were either experimentally intensified or relaxed through sex-limited selection. When antagonism was relaxed, alleles favoured under selection for female-typical larger size were consistently selected against in lines evolving male-typical smaller size, and vice versa. This pattern confirms their antagonistic effects. In contrast, intensified SA selection led to minimal genome-wide divergence, comparable to that observed under relaxed selection. This result is consistent with our previous pedigree analyses, which showed that SA lines maintained significantly more additive and dominance genetic variance in body size than lines under directional, sex-limited selection ^37^. Selection coefficient estimates further support this pattern. Responses in SA lines were more often aligned with male-optimal selection, and only rarely with selection for larger female size (Fig. 3b). This trend reflects the observed phenotypic divergence, particularly the lack of female size response under SA selection (Fig. 1) ^38^.

While these results demonstrate that SA selection can preserve genetic variation over short microevolutionary timescales, such as the 10 generations of artificial selection examined here, the minimal genomic divergence observed highlights the difficulty of detecting such selection using standard genome-scan approaches alone. This difficulty aligns with recent theoretical work suggesting that balancing selection, including that driven by SA trade-offs, may leave only weak or undetectable genomic signatures under realistic conditions, particularly when selection is weak or allele frequencies deviate from intermediate levels ^44^. Importantly, some loss of standing genetic variation was expected in our laboratory-based experiment, given the constraints of limited census and effective population sizes. Our aim was not to show complete preservation of variation, but to assess whether SA selection more effectively constrains allele-frequency change compared to directional, sex-limited selection under comparable demographic conditions. This focus on relative efficiency contrasts with interpretations that emphasise any variation loss under SA selection ^15^

Our second major finding concerns signatures of long-term constraint at the loci that responded strongly and divergently to sex-limited selection. These loci exhibit shifted allele-frequency spectra and elevated Tajima’s D values in the ancestral population to the selection lines, consistent with balancing selection. Prevalence of such polymorphisms may reflect both the frequency with which conditionally beneficial mutations arise and the range of mechanisms that can maintain them, including body size trade-offs ^45,46^. Until recently, the conditions for the long-term maintenance of balanced polymorphisms were thought to be restrictive ^47–49^. However, a variety of genetic and ecological mechanisms can broaden the conditions under which they are maintained. Among these is sex-specific beneficial dominance reversal, where an allele is dominant in the sex where it is beneficial but at least partially recessive in the sex where it is deleterious. This asymmetry can generate a net heterozygote advantage, which is a sufficient condition for balancing selection in diploid, randomly mating populations ^13,50,51^. A handful of empirical examples of sex-specific dominance have been documented ^27,52,53^, including in our study population Lomé ^38,54^. Sex-specific dominance variance in body size was maintained under the artificial SA selection while lost under sex-limited selection ^37^. Moreover, dominance variance in gene expression of a growth-regulatory pathway is sex-dependent in this population ^55^, providing a plausible mechanism for sex differences in phenotypic dominance. These findings call for a deeper investigation into the mechanisms by which dominance patterns may stabilize genetic variation in the context of balancing selection.

Our findings also have implications for understanding adaptive architecture, the set of loci that contribute to trait evolution. While the classic polygenic model predicts adaptation through small shifts at many loci with high redundancy ^30,36^, not all loci contribute equally, as effect sizes and levels of constraint can vary substantially ^30,36,56^. For example, pleiotropic loci may explain a large proportion of trait variation but contribute little to adaptation when selection is antagonistic ^57–61^. This was evident in our comparison between sex-limited selection for small male size and SA selection, both of which favoured smaller male body size. Divergence under sex-limited selection involved loci that responded weakly or differently under SA selection, despite the similar direction of selection and comparable male phenotypic responses (Fig. 1) ^38^. This highlights that the mode of selection, whether antagonistic or sex-limited, plays a key role in shaping adaptive genetic architecture, especially for traits with high intersex genetic correlation.

Finally, to complement our genome-wide insights, we examined the outlier loci for their involvement in biologically meaningful functions. Approximately a quarter of these SNPs are located within or near genes, most often in regulatory regions. Notably, eleven genes emerged as shared targets of positive selection for increased and decreased body size in the absence of SA trade-offs, with different SNPs within each gene concurrently responding to the opposite directions of selection. This pattern of antagonistic polyallelism is indeed predicted to evolve under SA selection in the presence of sex-specific dominance ^13^. A noteworthy polyallelic candidate locus is *Tolloid (Tld)*, a conserved member of the bone morphogenetic protein (BMP) signaling pathway. The flanking region of *Tld* shows elevated Tajima’s D in the ancestral population and responds strongly and directionally to size-limited selection. In contrast, allele-frequency changes at *Tld* under intensified sexual antagonism and relaxed selection are indistinguishable from one another and inconsistent between replicates. While essential for embryonic patterning and cellular growth ^62,63^, the role of BMP pathway in the development of adult body size is not fully understood, but has been repeatedly implicated ^64–70^ and demonstrated in the nematode *Caenorhabditis elegans* ^71^, as well as in the water strider *Microvelia longipes*, where it governs sexually selected leg elongation in addition to controlling body size ^72^. Altogether, this finding is consistent with the idea that balancing selection may preferentially act on conserved, developmentally constrained pathways where evolutionary flexibility is limited, a hypothesis that warrants further investigation.

## Conclusions

Using an experimental design that manipulates the strength of SA genetic trade-offs, we assessed both the short-term consequences and long-term genomic signatures of antagonistic selection on a complex life-history trait, body size. While genome scans showed that intensified SA selection helped maintain genetic variation, its effects on allele frequencies closely resembled those under relaxed selection, becoming apparent only in contrast to the strong, repeatable divergence observed under sex-limited selection for increased and decreased body size. Strikingly, the loci driving this divergence also carried signatures of long-term balancing selection in the ancestral population, suggesting that SA trade-offs may have preserved polymorphism over extended evolutionary timescales. Altogether, our findings demonstrate that SA selection can act not only as a constraint, maintaining genetic variation under opposing selection in males and females, but also as a source of adaptive potential, by preserving alleles that can respond rapidly once the trade-off is relaxed.

## Methods

### Study system

The cowpea weevil, *Callosobruchus maculatus*, is an important agricultural pest of legumes living in tropical and subtropical regions. Adult females lay eggs on the seeds of legumes where larvae develop. Under our laboratory conditions (at 29 °C, 50% relative humidity and a 12 h/12 h light/dark cycle, on *Vigna radiata*, the mung bean, as a host) development takes approximately 3 weeks, after which adults eclose and can complete their entire reproductive cycles without feeding or drinking. *C. maculatus* has 9 autosomes and a pair of sex chromosomes (XY sex determination system).

To test how different modes of sex-specific selection on body size affect genome divergence, we used lines subjected to 10 generations of replicated artificial selection, described in detail elsewhere ^38^. The lines were generated from an ancestral panmictic population, formed by pooling 41 isofemale lines established from females originally collected on an agricultural field near Lomé, Togo in 2010. Prior to establishing the selection lines, the ancestral population was kept under standard laboratory conditions for 10 generations, followed by a quantitative genetic pedigree experiment to describe genetic architecture of body size and sexual size dimorphism ^38^. The starting population for the selection lines, G0, was formed by the F2 generation of our breeding design ^73^. The selection lines were created by applying 10 generations of sex-limited (SL) family-level selection either only on males, towards smaller or larger size, only on females towards larger size, or sexually antagonistically (SA) on both sexes for larger females and smaller males (i.e., increasing the naturally occurring sexual size dimorphism). The lines also include a control subjected to relaxed selection on body size but otherwise experiencing similar demographic changes. Each mode of selection was independently replicated (replicate lines A and B). The male-limited selection treatments produced strong bi-directional divergence in male size. Females also evolved by correlational selection, but relatively less so due to a strong Y-linked variance affecting male size and subsequently sexual size dimorphism, which increased and decreased by ∼30% when selecting for smaller and larger males, respectively. Body size dimorphism did not change under drift alone (Relaxed selection), nor under female-limited selection due to similar increase in size of both sexes. It increased, however, by ∼50% under SA selection, due strong decrease in male size but no change in female size38.

### DNA extractions

Ancestral population and the artificial selection lines were sequenced using a Pool-seq approach at generation 6 and 10 (3 time points in total) ^74,75^, with each pool containing DNA of 50 males, resulting in 21 separate pools. Each sequenced pool was generated by pooling 10 extractions of 5 males in each. Males were randomly collected from the focal generations (0, 6, 10) and kept at-80°C. Prior to extractions the samples were sorted into groups of 5 size-matched males (weighed within a few hours from emergence) in 2 ml safe lock Eppendorf tubes. Pre-processing of all samples started with homogenization with TissueLyser II (15 Hz, 2 x 20s) in ATL Buffer (Qiagen DNeasy Blood and Tissue kit) and with two 5 mm stainless steel beads in each Eppendorf. After mechanical disruption, samples were incubated overnight with 20 µl of proteinase K (20 mg/ml) in a thermomixer (400 rpm, 56°C). The rest of the procedure followed the Qiagen DNeasy Blood & Tissue kit protocol. DNA was eluted in EB (EDTA-free elution buffer) by passing eluate twice through the column. The quality of each sample was assessed with Nanodrop 2000 and Qubit 2.0 (using DNA BR kit). Qubit DNA concentration readings were used to evenly pool DNA from 10 samples, each consisting of DNA extracted from 5 males, ensuring an equal contribution of each sample in the final pool. For each sample, several pools were prepared, and the sample of the highest quality was selected for sequencing.

### Resequencing

Libraries for each of the 21 pools were prepared using TruSeq PCRfree DNA sample preparation kit with a targeted insert size of 350bp. Libraries were then sequenced on Illumina NovaSeq 6000, paired-end 150bp read length (S4 flow cell and v. 1.5 sequencing chemistry). Ancestral population was sequenced on a single lane, while all other selection line pools were randomized and sequenced across 4 lanes (one S4 flow cell).

### Mapping pipeline

The mapping pipeline was implemented using Snakemake 7.6.2 ^76^. Because the ancestral population and the selection lines differed in sequencing design and computational requirements (see below), they were processed separately. First, all reads were trimmed using cutadapt 4.0 ^77^. We removed primers specific to TruSeq libraries, enforced a base quality score of 20 and a minimum read length of 75. After inspecting quality of trimmed reads with FastQC 0.11.9 ^78^ we mapped them to the *C. maculatus* reference genome assembly (originating from the same population, from Lomé, Togo, ENA ID SAMEA113578947 ^39^, using BWA-MEM 7.17 ^79^. The ancestral population was sequenced at substantially higher coverage, which produced much larger data volumes, and these reads were therefore mapped using the faster BWA-MEM 2 v. 2.2.1 ^80^. Filtering was conducted with SAMtools v. 1.14 ^81^. We retained reads of quality score above 20 and excluded secondary alignments (-F 0×100 flag). For the selection lines, whose libraries were distributed across four sequencing lanes, we received four lane-specific FASTQ files per line and merged the resulting BAM files after mapping using SAMtools v. 1.14 ^81^. Duplicates were removed with Picard MarkDuplicates v. 2.23.4 ^82^ and realignment around indels performed using GATK v. 3.8 ^83,84^. Statistics for bam files were collected at several time points using Qualimap v. 2.2.2d ^85^ and SAMtools v. 1.1481.

### SNP calling and creation of the sync file

We performed variant calling using the ancestral population of the selection lines. The identified sites served as a reference for subsequent analyses of our selection lines, based on the assumption that adaptation in these lines proceeded from standing genetic variation present in the ancestral population. To ensure accurate identification of variants, we employed two different variant callers. Firstly, we utilized PoolSNP v. 1.05, a heuristic variant caller specifically designed for variant detection in pooled samples ^86^. Secondly, we applied freeBayes ^87^, which, while not designed for Pool-seq data, outperforms GATK when used for variant calling of Pool-seq data ^88^. For our purposes, freeBayes was run as implemented in Grenepipe v. 0.10.0 ^88^, a comprehensive pipeline that supports Pool-seq applications. Variant calling with freeBayes was followed by GATK variant filtration implemented in the same pipeline with default settings; we excluded SNPs with a quality by depth (QD) less than 2.0, indicating insufficient data per allele, and those showing a Fisher strand (FS) score greater than 60.0 to eliminate potential sequencing or alignment artifacts due to strand bias. Additionally, SNPs where the average mapping quality (MQ) of supporting reads was below 40 were removed. While further processing both VCF files with bcftools v. 1.14 ^81^, we removed multiallelic sites in both datasets and excluded sites overlapping RepeatMasker-annotated regions ^89^ from the FreeBayes VCF using a curated repeat library for *C. maculatus* ^90^. The latter step resulted in a removal of about 70% of sites; indeed *C. maculatus* is known for having a very high content of repetitive sequences, consisting mainly of DNA transposons followed by LINE retrotransposons ^90^. Finally, two pre-filtered VCF files were intersected using bcftools v. 1.14 (isec-p), yielding 9 225 886 overlapping variants (out of 12 678 677 sites from freeBayes and 18 887 140 from PoolSNP). We then created Pool-seq specific synchronized files (sync files), which list base counts at all genomic positions ^91^, using Grenedalf v. 0.2.0 ^88^. Specifically, individual sync files were created for each selection treatment, including the ancestral population, and two replicates from each experimental treatment at generations 6 and 10. This was achieved by processing pileup files for all populations, including the ancestral population. To ensure data quality, the minimum base quality was raised to 30 (--pileup-min-base-qual 30) ^88^. Additionally, during the creation of these sync files, we limited our analysis to sites present in the final VCF file (--filter-region-vcf). Subsequently, a comprehensive final sync file was compiled using Grenedalf v. 0.2.0 by consolidating data from all four treatments. Further filtering was conducted using R 4.2.2 ^92^ and included: 1) removing sites fixed for the alternative allele in the ancestral population, so that only sites that were segregating in the ancestral population were preserved, 2) keeping only biallelic SNPs with the same alleles as in the ancestral population throughout the entire dataset, 3) excluding contigs shorter than 100 kb ^39^, and 4) filtering based on coverage of minimum 10 (for all generations and treatments) and maximum of 95^th^ percentile (554; only based on the coverage for the ancestral population). Such a processed sync file contained 5 398 991 highest quality genomic sites. Finally, using the R package poolSeq we created a final file, where allele frequencies across selection treatments and generations were polarized for the ancestrally minor allele ^43^. This file served as a basis for our analyses.

### Population genetic analyses

To examine genetic differentiation and population structure among derived artificial selection populations, we used the R package pcadapt v. 4 ^42^, which provides biplots for visualizing population clustering and PCA-based selection scans to identify outliers. Specifically, we employed the component-wise selection scan, which identifies SNP outliers along a particular PC axis based on SNP loadings. The PCA-based outlier approach allows focusing on the axis of variance that recapitulates the selection history. Moreover, this approach was selected for its capacity to handle large datasets, such as those with multiple time points and selection regimes, as in our artificial selection experiment. The pcadapt package further facilitates analysis by supporting the direct use of Pool-seq frequency files.

To complement the approach implemented in pcadapt, we also used AF-vapeR ^41^. SNPs were split into non-overlapping windows of 20 SNPs, and within each window we computed allele-frequency change vectors for predefined contrasts. Using a common-ancestor design, we constructed vectors from the ancestral population to generation 10 for each treatment × replicate. AF-vapeR then performed an eigen-decomposition of the covariance matrix of these change vectors, with large eigenvalue 1 (EV1) indicating coordinated change (parallel or anti-parallel).

For subsequent analysis of genetic differentiation between populations, we chose the unbiased Nei estimator of *F_ST_*, as implemented in Grenedalf v. 0.3.0 ^88^. For *F_ST_* and PCA-based analyses, we set the frequency threshold for the ancestrally minor allele to 0.05. We assessed genetic diversity levels in the regions of outlier SNPs by calculating genetic diversity metrices, π and θ Watterson, along with Tajima’s D. These calculations were performed using Grenedalf v. 0.3.0 ^88^ directly on the final sync file.

Selection coefficients (*s*) were estimated with estimateSH from the poolSeq package^43^ as the slope of a least-squares fit to logit-transformed allele frequencies between generations 0 and 10, assuming codominance (h = 0.5, which was used as a standard approximation in the absence of locus-specific dominance information). Replicate-specific N_e_values were estimated separately by the same package and supplied to estimateSH. Positive values of *s* indicate increasing allele frequency and negative values decreasing frequency over the interval. Rare extreme estimates (|*s*| > 1) occurred at a very low rate (≈ 10^-4^ of sites) and were interpreted as overestimation in cases of rapid allele-frequency shifts over a short timescale combined with small N_e_ (below 20 in our case).

### Functional characterisation of the outlier loci

Outlier SNPs were first mapped to nearby genes by identifying variants located within 2 kb upstream or downstream of gene boundaries. Gene Ontology (GO) enrichment analysis of the resulting gene set was conducted using topGO ^93^. Putative *Drosophila* orthologs of the genes with SNPs responding to both directions of SL selection were identified using BLASTP with default parameters.

All intermediate data manipulations and plotting were conducted using R 4.2.2 ^92^ with packages from the tidyverse collection, including ggplot2 ^93^. For efficient data handling, we also utilized the data.table package ^94^

## Acknowledgements

We thank Arild Husby and David Berger for their helpful and insightful comments. Sequencing was performed by the SNP&SEQ Platform at Uppsala University. The facility is part of the National Genomics Infrastructure (NGI) Sweden and Science for Life Laboratory. The computations were enabled by the Swedish National Infrastructure for Computing at UPPMAX. The work was supported by grants from the Swedish Research Council (Vetenskapsrådet) (no. 2019-05038 and 2023-04869), and Carl Tryggers Stiftelse to E.I. (grants no. CTS-18:163 and CTS-19:155), and the Birgitta Sintring Foundation to M.K.Z. (grant no. S2024-0007).

