## Supplementary Figures for "From constraint to opportunity: Relaxing sexual antagonism reveals adaptive potential maintained by balancing selection"

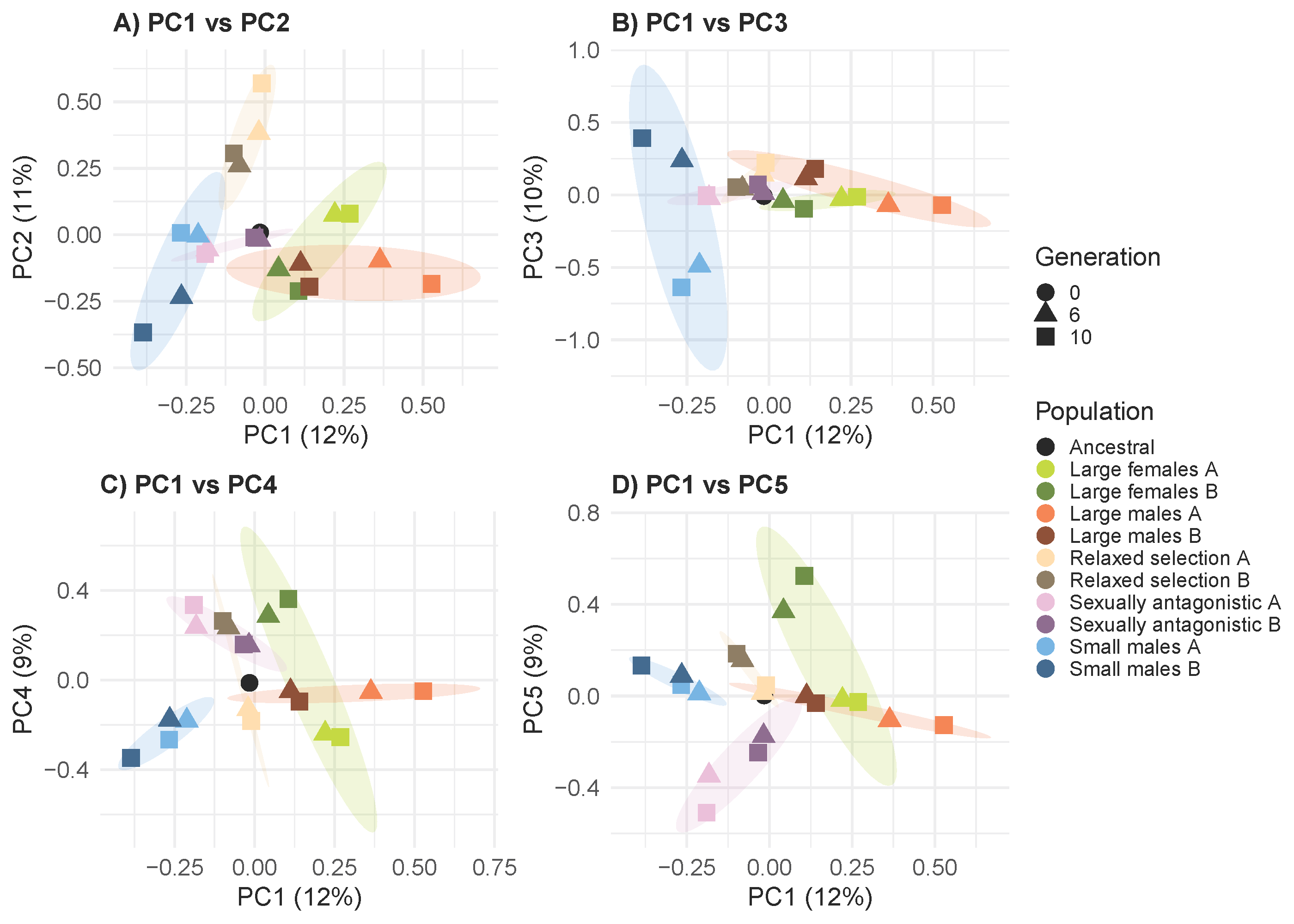

**Figure S1**. PCA biplot on SNP frequencies in the autosomal loci across the first 5 PCs.

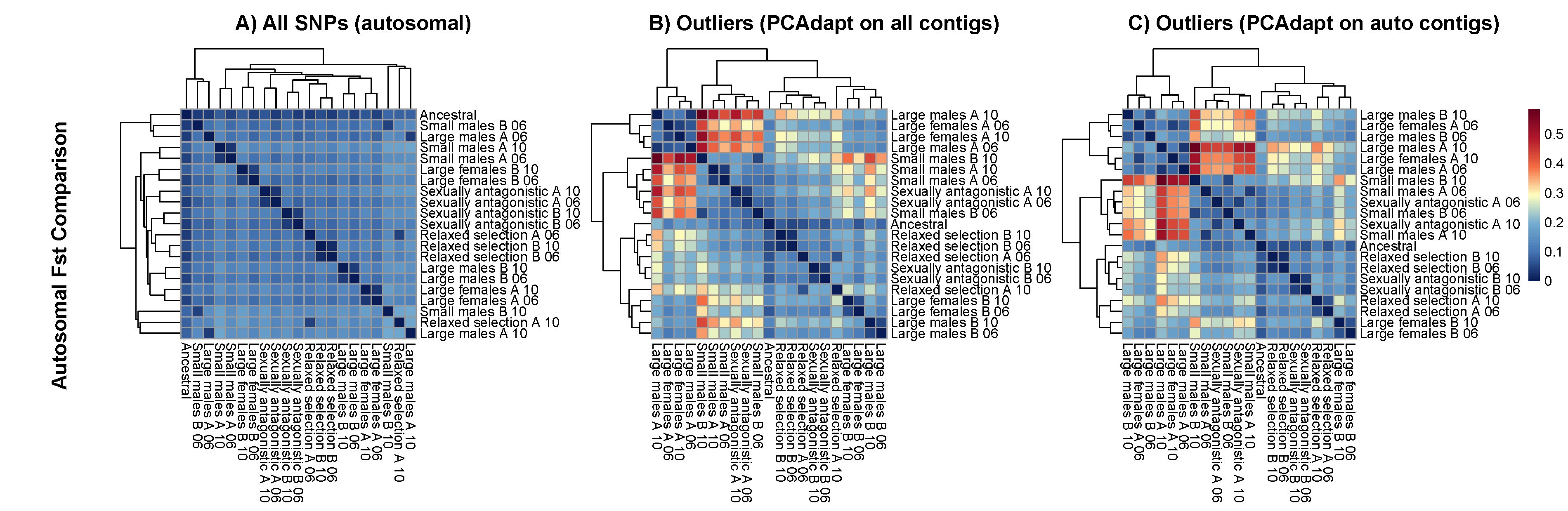

**Figure S2**. F_ST_ pairwise correlations for a) all autosomal SNPs, b) all outlier SNPs, c) autosomal outlier SNPs. In (a) All populations remain closest to the ancestral population, while divergence from the ancestor increases from generation 6 to 10. In (b) and (c), higher and more structured F_ST_ values reveal increased differentiation at outlier loci, particularly between sex-limited selection lines selected for large vs. small body size, highlighting the effects of divergent selection.

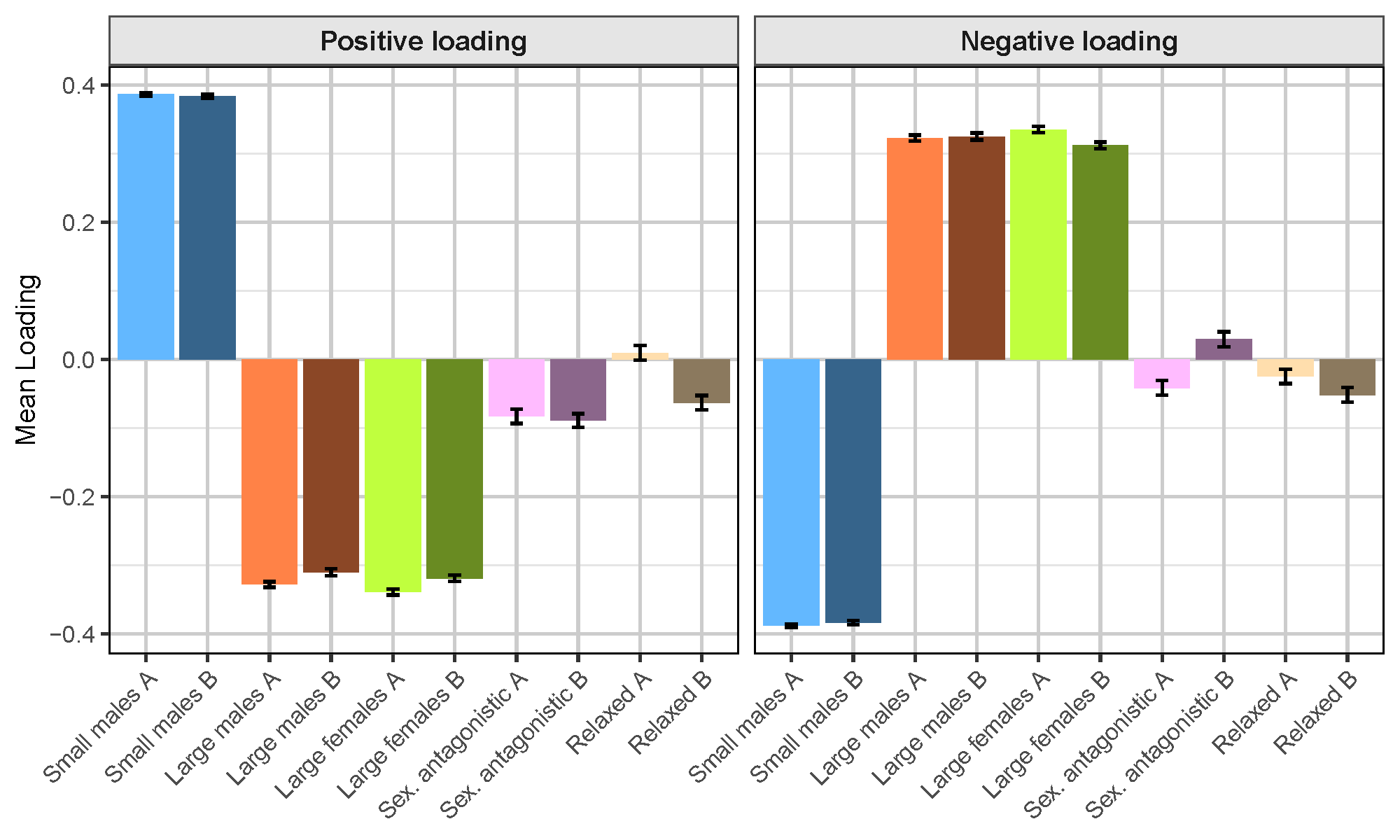

**Figure S3**. Mean eigenvector 1 loadings across all lines for the 100 windows that maximize divergence between the bidirectional sex-limited selection lines (i.e. SL small male selection vs. SL large female *and* SL large male selection).

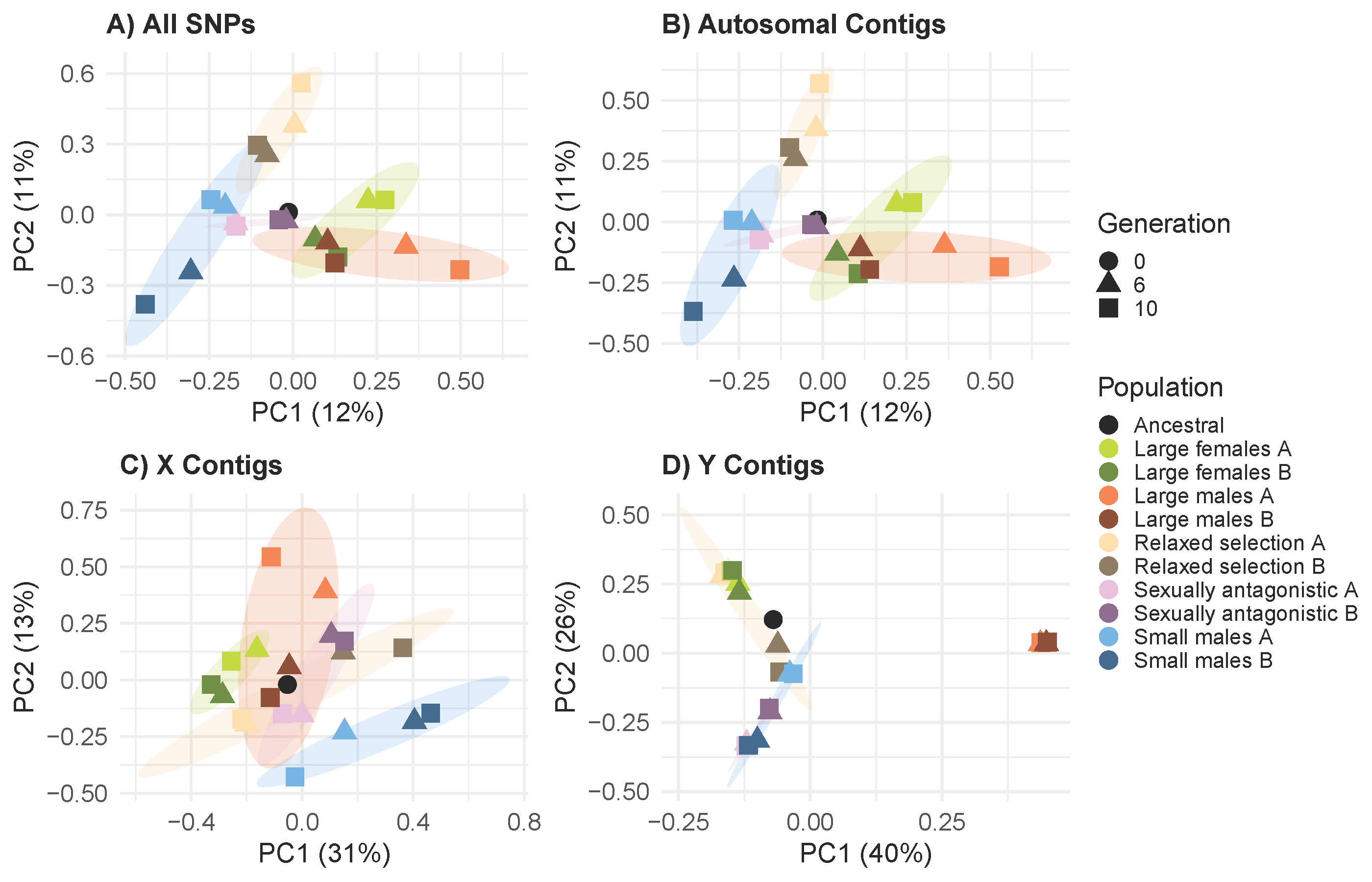

**Figure S4**. PCA biplots of the first two PCs for a) all, b) autosomal, c) X-linked, d) and Y-linked SNPs.

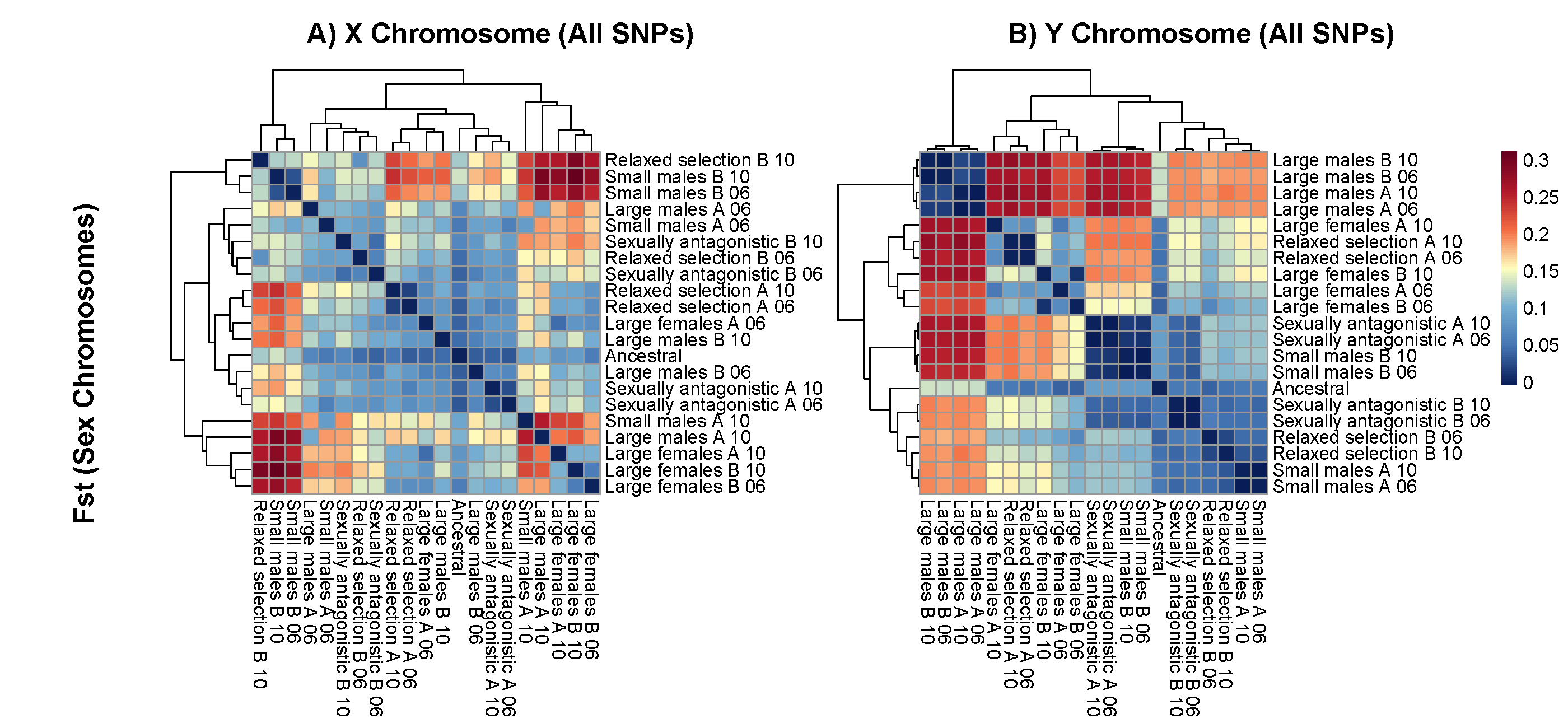

**Figure S5**. F_ST_ pairwise correlations for a) X -linked and b) Y-linked SNPs.

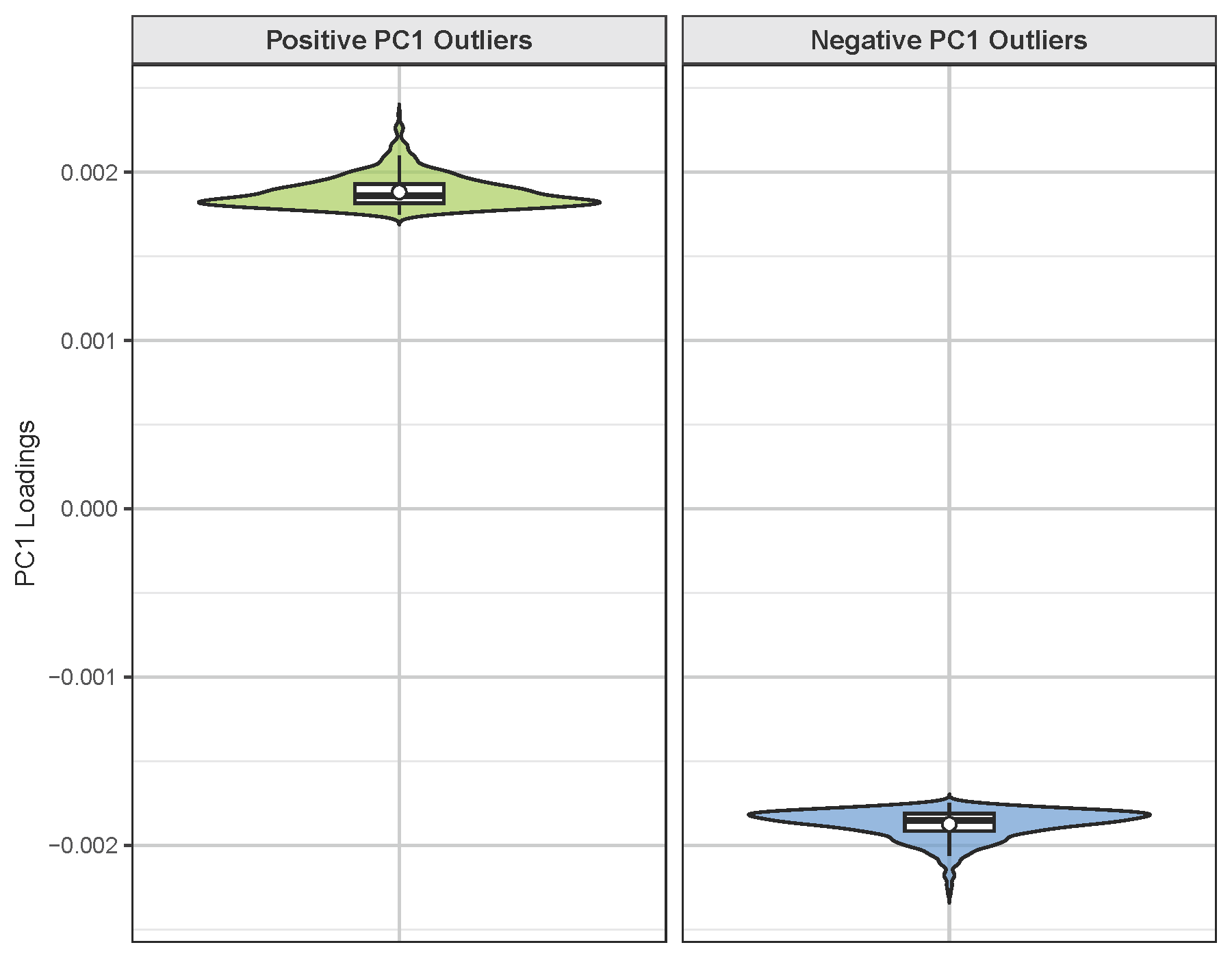

**Figure S6**. Distribution of PC1 loadings for SNPs identified as significant outliers (q < 0.05).

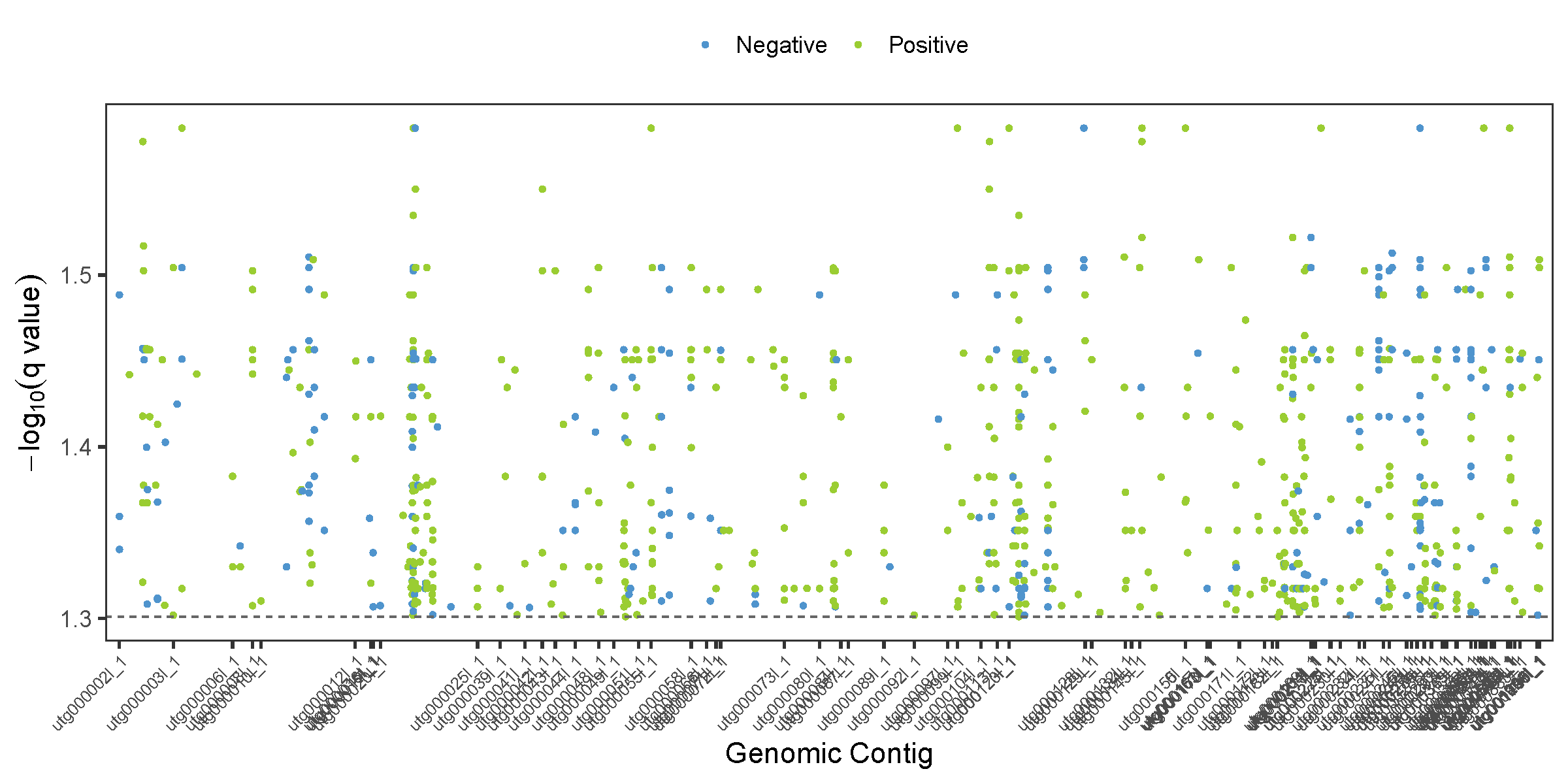

**Figure S7**. Manhattan plot of the outlier SNPs across the contigs.

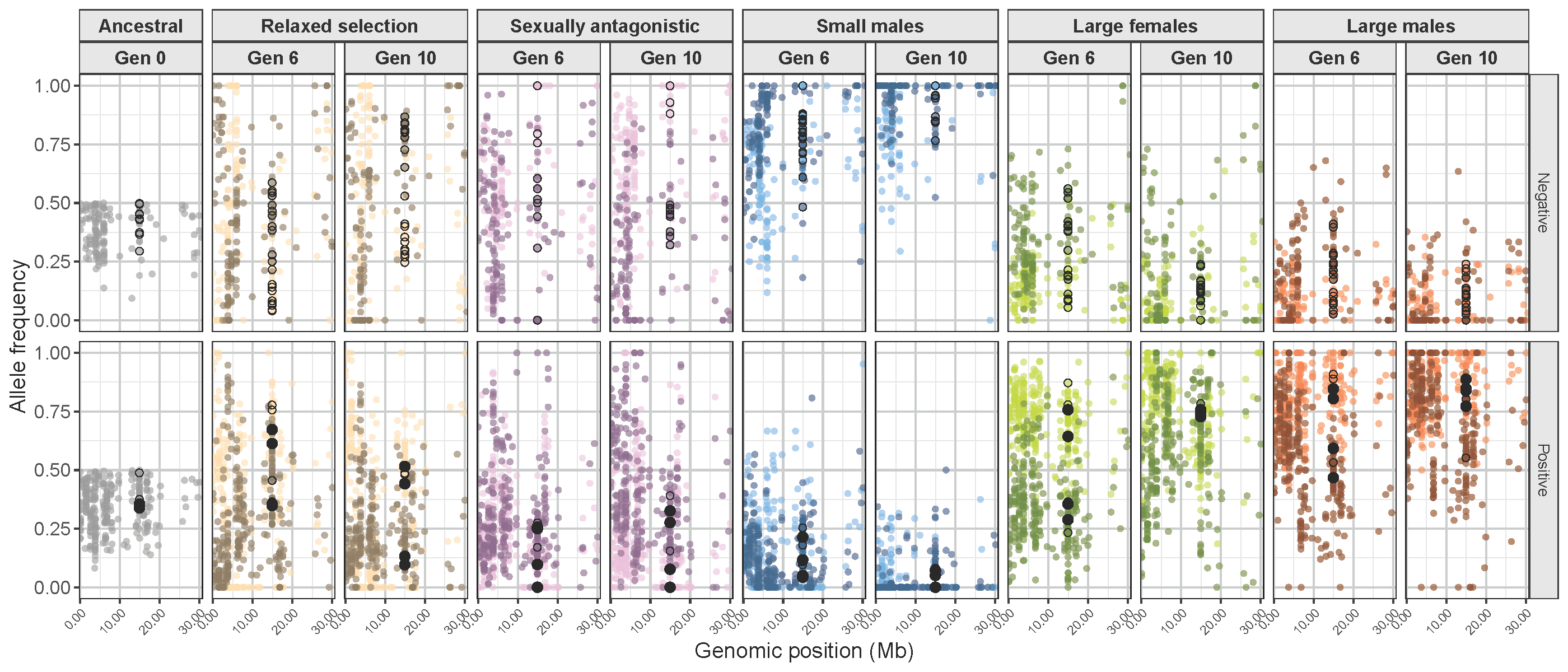

**Figure S8**. Allele frequency changes in the *Tolloid* locus and its flanking regions. Genomic sites identified as outliers (with the q value of 0.10) are marked with black rings. Regions within the gene body are additionally filled.
